# *Hashi*: Bridging Statistical Model Derived 1D Microstate Encodings and Protein 3D Structural Ensembles

**DOI:** 10.64898/2026.06.01.729173

**Authors:** Harshavardhan Madhan, Athi N. Naganathan

## Abstract

The functioning of proteins is intimately linked to the conformational states they sample within the native ensemble. Generating ensembles from a single static structure is therefore a research domain receiving considerable attention. In this application note, we introduce *Hashi*, a pipeline to rapidly generate realistic structural ensembles from the outputs of the structure-based Wako-Saitô-Muñoz-Eaton (WSME) statistical mechanical model of protein folding. This approach relies on integrating the block WSME model outputs – strings of zeros and ones describing the conformational status of every residue over thousands or millions of microstates each assigned a statistical weight derived from physically grounded energy-entropy terms, and free energy profiles - with the RANCH module of the EOM (ensemble optimization method) from the ATSAS software suite, providing three-dimensional views of the structural ensembles within the model framework. It is applicable to a variety of single-chain monomeric systems with lengths ranging from 30 to 500 residues, including globular and repeat proteins. Ensembles can be generated within seconds to minutes on a standard computer, without recourse to high-performance computational clusters. The generated structural ensembles can also be rank-ordered according to their free energies within a given macrostate or a range of reaction coordinate values. Since the statistical weights of the WSME model microstates can be reweighted or calibrated with experiments, the ensembles shed light on not just the folding mechanism but also on the structural excursions that determine function and opening of otherwise buried binding pockets.

## Introduction

Generating protein conformational ensembles given a static three-dimensional (3D) structure is one of the longstanding goals of computational techniques. This is because the functioning of proteins is not determined by a single structure but an array of structural excursions^1–4^ whose relative probabilities define opening-closing events involving structural pockets, melting of helices or even large-scale inter-domain motions. The ideal way to achieve this objective involves accumulating conformations as a function of time in all-atom^5–8^ or coarse-grained MD simulations^9,10^ or through enhanced sampling techniques,^11–13^ identifying regions of proteins that display enhanced flexibility, and projecting these conformations on to specific coordinates or order parameters to arrive at near-equilibrium distributions. Recently, generative AI methods trained on data-rich MD simulations have upended the field by generating physically and thermodynamically plausible state distributions,^14–18^ establishing powerful alternatives to conventional MD methodologies.

The structure-based^19^ Wako-Saitô-Muñoz Eaton (WSME) statistical mechanical model of protein folding^20,21^ occupies a sweet spot between these two extremes. In its original construction,^20^ the WSME model represents each protein residue using a binary conformational variable, taking the values *1* or *0* to denote the folded and unfolded status, respectively. This leads to 2*^N^* microstates for a *N*-residue protein, wherein every microstate is represented as a string of *ones* and *zeros*. Multiple approximations^21,22^ and versions^23–26^ have been developed over the last three decades on this model. Importantly, despite being phenomenological and Gō-like,^19^ the WSME model framework has been adept in capturing the folding mechanisms of numerous proteins,^27–33^ engineer protein stabilities,^34^ understand long-range mutational effects,^35,36^ map the thermodynamic and evolutionary design principles of protein structures^36–38^ and to quantify folding pathway heterogeneity.^39,40^ The energy-entropy function is physically sound, with the parameters derived from experiments directly,^41^ empirically^23^ or through MD simulation forcefields.^26^ The rapidity of the method is an added advantage, with the total partition function calculated within seconds to minutes for most proteins, using either transfer-matrix formalisms^20,42^ or algorithmic enumerations.^43^

While the mean folding probability of every residue at different points along the reaction coordinate are readily obtained from the WSME model, a 3D view is not currently possible. In this application note, we bridge this divide through the *Hashi* (*bridge* in Japanese) workflow. We integrate the WSME model outputs – particularly, the strings of *ones* and *zeros* – with the RANCH module of the highly successful ensemble optimization method (EOM) from the ATSAS software suite,^44,45^ which is typically used for predicting SAXS profiles. The module reads in the folded segments (islands of *ones*) and the sequence of the disordered linkers (islands of *zeros*) and rapidly provides a structural ensemble that is consistent with input constraints. The ensembles generated can be employed to understand folding mechanisms and even the functioning of proteins, especially those that require large structural motions.

## Methods

### bWSME model

The description of the block WSME (bWSME) model, its intrinsic approximations and validation, have been detailed elsewhere,^22,36^ and we summarize its key aspects here. The bWSME model that we use in this work considers stretches of consecutive residues, termed blocks, as the fundamental folding units. If three consecutive residues are considered as a single block, this leads to 2*^n_b_^* microstates where *n_b_* = *N*⁄3 (*N* being the number of residues in the protein). Blocks are not allowed to span secondary structure boundaries and hence a block length of 3 effectively means that many of the blocks (and not all) have a length of 3. In addition, the current approach considers microstates derived from three sequence approximations – single and double sequence approximations (SSA and DSA, respectively) that account for single stretches and two stretches of folded residues/blocks separated by *zeros* (unfolded regions), and DSA with loop (DSAw/L) that allows for interactions across the folded islands despite the intervening *zeros*. This further reduces the number of microstates to the binomial coefficients – 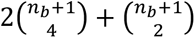 - apart from the fully unfolded microstate (a string of only *zeros*). If the protein is small enough, like Villin (*N* = *n_b_* = 35), a residue level treatment can be considered as it leads to only 118,440 microstates. For a large protein like Pertactin with 539 residues, a block size of 5 leads to *n_b_* = 137 and hence the number of microstates within the model framework will be 28,935,633.

The statistical weight *w_i_* of microstate *i* (the strings of *ones* and *zeros*) is calculated from a (free)energy-entropy function, Δ*G_i_* = Δ*G^stab^* − *T*Δ*S_conf_*. The stabilization free energy Δ*G^stab^* includes contributions from native van der Waals interactions (*E_vd_*_W_), electrostatics (*E_elec_*) and simplified solvation (Δ*G_solv_*). The van der Waals interactions are identified with a 5 Å (typically) heavy-atom distance cutoff from the input structure, while the electrostatic interactions are accounted for by a Debye-Hückel formalism with all-to-all electrostatics.^24^ The solvation free energy term scales with the number of interactions within a given microstate and the heat capacity change per native contact (Δ*C_p_^cont^*) together with the associated temperature dependence of solvation free energy. The model invokes a large entropic penalty for fixing a residue in the native conformation and this Δ*S_conf_* (units of J mol^-1^ K^-1^ per residue) can be fixed for all residues or modulated according to the secondary-structure status. In other words, residues identified as coil or glycine are assigned a larger entropy penalty of Δ*S_conf_* + ΔΔ*S* where ΔΔ*S* = −6.06 J mol^-1^ K^-1^ per residue,^25^ and proline a zero entropic penalty given its limited flexibility.

The total partition function is calculated as *Z* = ∑*_i_ w_i_*, from which free energy profiles are generated as a function of number of structured blocks or residues, the natural reaction coordinate of the model. While microstates correspond to individual binary strings, a macrostate is defined by grouping together all microstates with the same number of structured blocks (*ones* in the binary string representation). One of the outputs of the model is the matrix *pepval* that has 7 columns with the rows corresponding to the microstates, and the columns providing the information on the number of structured residues/blocks, statistical weights, and positions of folded and unfolded residues (Figure 1, Supporting Figure S1).

**Figure 1.**
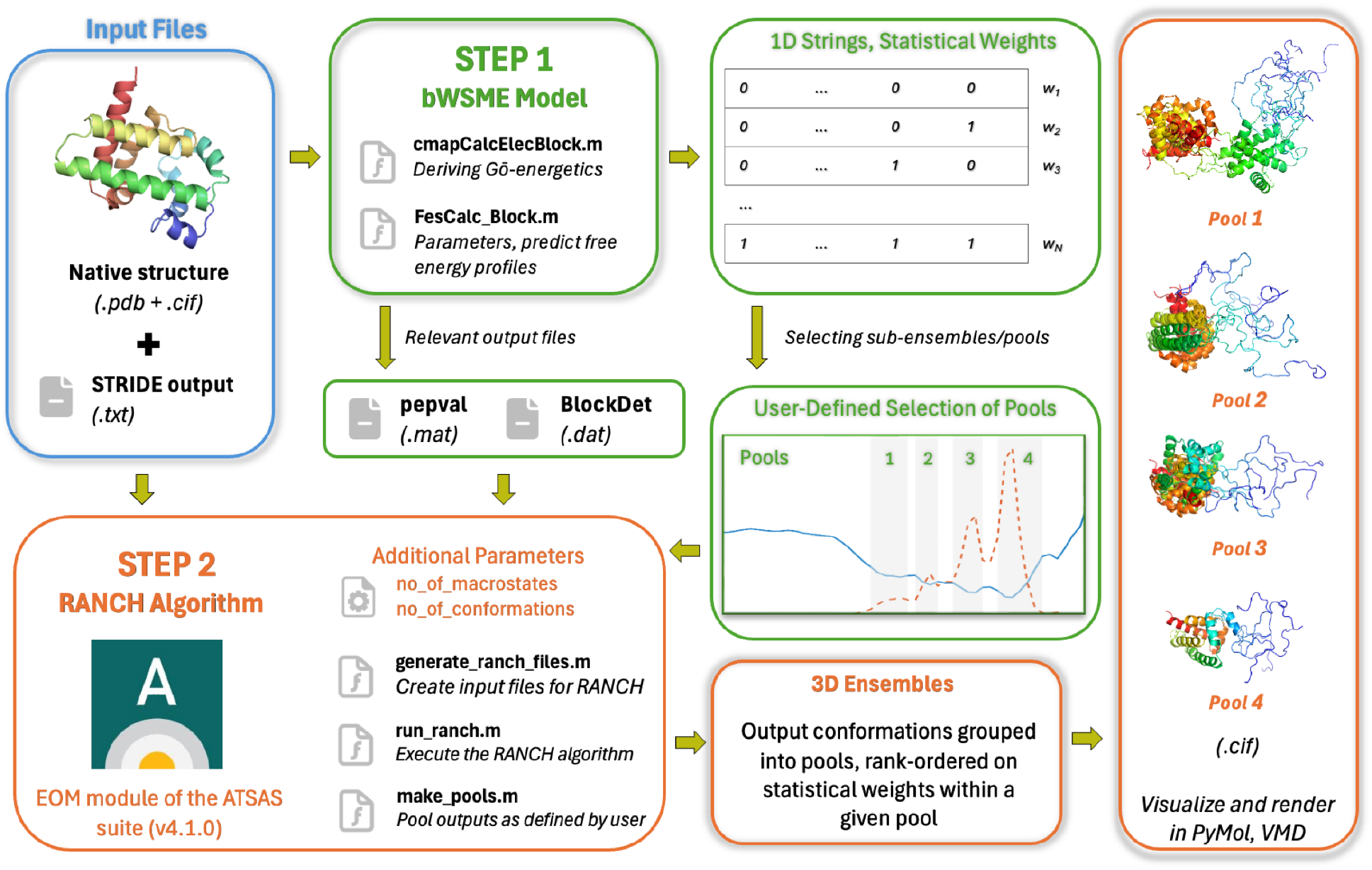
Overview of the *Hashi* workflow for a sample protein. Input file, highlighted by the blue box (.pdb and .cif structures, along with formatted STRIDE output), are processed in two steps. In the first step (Step 1 – bWSME Model – highlighted by green boxes), the files are processed to derive Gō-energetics from the input structures, free-energy profiles are generated from the bWSME model, and sub-ensembles (*macrostate_pools*; i.e. the shaded regions in the free energy profile) are manually set by the user. In the second step (Step 2 – RANCH Algorithm – highlighted by orange boxes), 3D ensembles are generated for these pools, using the output files from the bWSME model as its input.

### Generating Ensembles

The step-by-step procedure to generate the *pepval* matrix and free energy profiles is provided in the supporting text and the associated Github page: https://github.com/AthiNaganathan/Hashi. Running the script main_WSME.m generates both the 1D free energy profile and the *pepval* matrix in about a *second* for Villin, and in *∼15 minutes* for Pertactin (block lengths 1 and 5, respectively), on an Apple M3 or Intel i5 processor with 16 GB RAM. For a fixed entropic penalty value, strengthening the vdW interaction energy (i.e. making it more negative) will stabilize the folded state relative to the unfolded state. For a protein with a size between these two extremes (i.e. Villin and Pertactin), the bWSME model outputs can be generated within a few seconds to minutes.

Once the *pepval* matrix is defined, it is straightforward to generate 3D ensembles using the RANCH module of EOM^44,45^ in ATSAS.^46^ This requires a specific set of files within the working directory: the *pepval* matrix, a *Blockdet* file (containing block-to-residue mapping information), a PDB file, and a CIF (Crystallographic Information File) (Figure 1). These files are automatically generated at the end of running the bWSME model, but the user may also provide them manually, if available from a previous run. The user then defines the range of reaction-coordinate values over which the ensemble (or pool) is generated, specified by the variable *macrostate_pools = [x y]*, where *x* and *y* correspond to the shaded regions of the free energy profile (Figure 1). Two additional variables are then required: the number of rank-ordered microstates sampled from within this pool (*no_of_microstates*) and the number of conformations generated per microstate (*no_of_conformations*). Because an unfolded segment is represented as a string of zeros, a given microstate is consistent with multiple 3D structural representations; the *no_of_conformations* variable accounts for this degeneracy. These are generated in the RANCH module by considering Cα–Cα Ramachandran statistics at a single Cα per unfolded residue while avoiding steric clashes. The generated structural representations are rank-ordered and carry information on the number of structured blocks/residues from which they were chosen, the bWSME model approximation in which they fall (column 7 of *pepval* matrix) and the conformation number in their file name (Figure 1, S1). The *CIF* files of the different conformations generated in the folder can be colored, superimposed with the starting structure and used for additional analyses. For the disordered regions, mainchain and sidechain atoms can be added through standard tools, including PULCHRA^47^ or SCWRL.^48^

## Results and Discussion

### Conformational ensemble of Villin and the folding mechanism of Pertactin

The Villin headpiece, a small 35-residue helical protein (Figure 2A), has served as a prototypical model system in protein folding studies.^39,49–53^ The predicted 1D free energy profile of Villin matches with the WSME model employed by Eaton and coworkers,^39^ exhibiting a broad native well centered around 30 structured residues and a shallow intermediate-like state at ∼15 structured residues (Figure 2B). The broad native well can arise from either of the first or the third helices undergoing large-scale fluctuations. The 2D surface, constructed with the coordinates being the number of structured residues at the N- and C-terminal halves of the protein, shows that it is the C-terminal half of the protein that exhibits a broader minimum with the possibility of up till 6 residues unfolded (Figure 2C). The generated native ensemble of Villin through the *Hashi* workflow matches with this expectation with large unfolding of the C-terminal half of the third helix (Figure 2D). Our predictions agree with the experimental results of Kiefhaber and coworkers, who employed triplet-triplet energy transfer studies to show that the third helix undergoes a rapid ‘unlocking’ reaction.^54^

**Figure 2.**
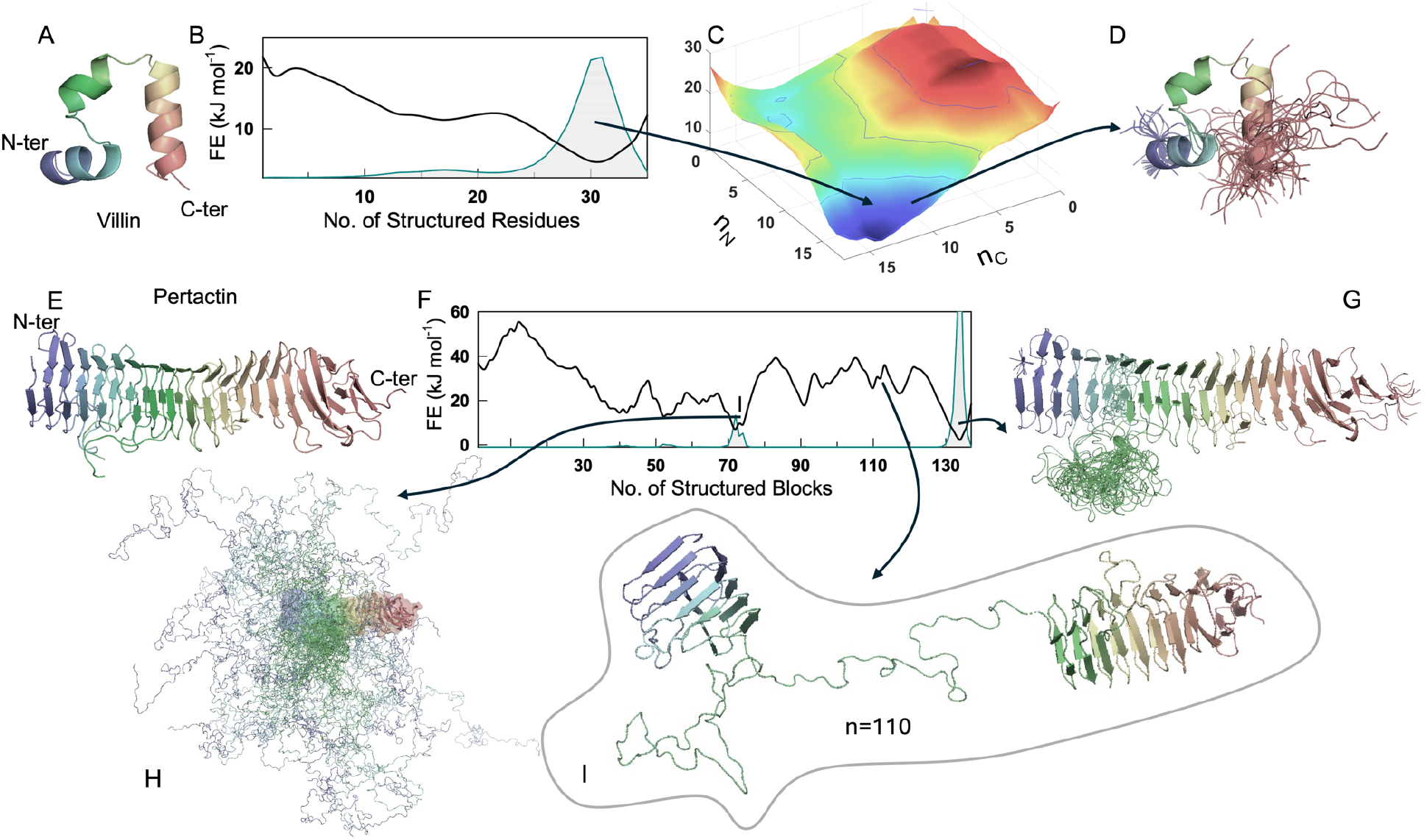
Conformational ensembles of Villin and Pertactin. (A) Structure of Villin (PDB id 1VII). (B) One-dimensional free energy profile of Villin as a function of the number of structured residues, the reaction coordinate (RC). The gray shaded area represents the probability density. (C) A two-dimensional conformational landscape as a function of the number of structured residues in the N- and C-terminal half of the protein (nN and nC), with the z-scale in free energy units of kJ mol^-1^. (D) Generated native ensemble from the *Hashi* workflow colored in the spectral scale from N- to C-terminus (blue to red) for the following user-defined parameters: *macrostate_pools = [25 34], no_of_microstates = 20, no_of_conformations = 5*. (E) Structure of Pertactin (PDB id: 1DAB). (F) Same as panel B, but for Pertactin. (G, H, I) Generated ensembles for the folded state (panel G), a low free energy intermediate state (panel H) and high free energy intermediate (at a RC value of 110; panel I). The parameters are: *macrostate_pools = [128 137; 107 111; 68 76], no_of_microstates = 5, no_of_conformations = 10*.

Pertactin, a large 539 amino-acid monomeric β-rich protein (Figure 2E), has been shown to fold *via* an intermediate in which only the C-terminal half of the protein is folded.^55^ The bWSME predicted free energy profile of Pertactin is flat with multiple shallow barriers and peaks (Figure 2F). The folded well is quite sharp with minimal width. Accordingly, the predicted 3D ensemble from the *Hashi* workflow showcases a well-folded system with only the long loop between residues 225-263 displaying multiple conformational states (Figure 2G). A stable intermediate *I* is visible at ∼72 structured blocks, and this corresponds to disorder in the N-terminal stretch of 1- 263 residues (i.e. the N-terminal half), with a folded C-terminal half (Figure 2H). The Pertactin variant spanning residues 1–334 is fully disordered both in isolation and in the intermediate state of the full-length protein,^55^ mirroring our predictions. The unfolding mechanism is better observed at a reaction coordinate value of 110 (i.e. when 110 blocks are structured). The entropic stability of the loop dominates the landscape leading to the separation of the N-terminal and C-terminal regions (Figure 2I), which serves as a precursor to the more stable intermediate with the disordered N-terminal region. Note that the snapshot shown is just one possible 3D representation that is consistent with the conformational status of residues derived from the microstate strings.

### Experimental validation and generation of structural ensembles for a database of proteins

To assess if the generated structural ensembles capture experimental trends, we selected three model systems – ACBP, TipAS, and I*κ*B*a*. The C*a*-RMSF profile of the *Hashi*-derived 3D ensemble agrees well with the regions that exhibit maximal positional fluctuations in all-atom MD simulations of ACBP (Figure S2A-S2D).^56^ Similarly, for the antibiotic-sequestering protein TipAS,^57^ the *Hashi*-derived ensemble reproduces the experimental SAXS profile (through the CRYSOL module of ATSAS) substantially better than the PDB structure, consistent with the disordered N-terminal domain observed experimentally (Figure S2E-S2H).^57^ Finally, for the repeat protein I*κ*B*a*, the experimental hydrogen-deuterium exchange mass spectrometry (HDX-MS) data^58^ strongly correlates with the RMSD of the individual repeats relative to the static structure, highlighting protein regions that undergo enhanced structural excursions (Figure S2I-S2L). It is, in fact, possible to tune the contact map or the associated energetics of interactions in the bWSME model to better capture experimental trends including HDX-MS profiles,^59^ heat capacity curves^29,56,57,59,60^ and mutational effects.^41^ Close agreement with experiment is therefore readily achievable with the bWSME model, and the *Hashi* pipeline now provides a structural view of the resulting ensemble (*vide infra*).

We further compiled a small database of 15 proteins spanning a range of sizes (65–375 residues), topologies, and secondary structure compositions. For each protein, native ensembles comprising the top 20 microstates (ranked by statistical weight) and 5 conformations per microstate can be generated in 5–20 seconds on an Apple M3 or an Intel i5 processor with 16 GB RAM (Figure 3), once the *pepval* matrix has been computed. We then performed extensive *Hashi* runs for four large proteins – TipAS (143 residues), Kemp Eliminase (299 residues), p38 (355 residues) and Pertactin (535 residues) – generating ensembles along the reaction coordinate (see Supporting Text and Figure S3). While no systematic trend emerged, larger proteins required marginally longer generation times with 100 conformations generated within 40 seconds on average with little dependence on the degree of disorder – that is, conformations could be generated rapidly regardless of how many residues were disordered. However, there were two notable exceptions. The RANCH module took ∼3 minutes to generate conformations for certain microstates in p38; additionally, it failed to generate conformations for certain microstates in Pertactin with disorder in specific sequence stretches (SI text and Figure S3, S4). This could be a consequence of the tight turns in this repeat protein that disallowed the generation of multiple alternate conformations. In the current version of *Hashi*, attempts are made to generate conformations for two minutes before being timed out; these microstates are skipped and a warning message is provided to the user including the identity of the microstates.

**Figure 3.**
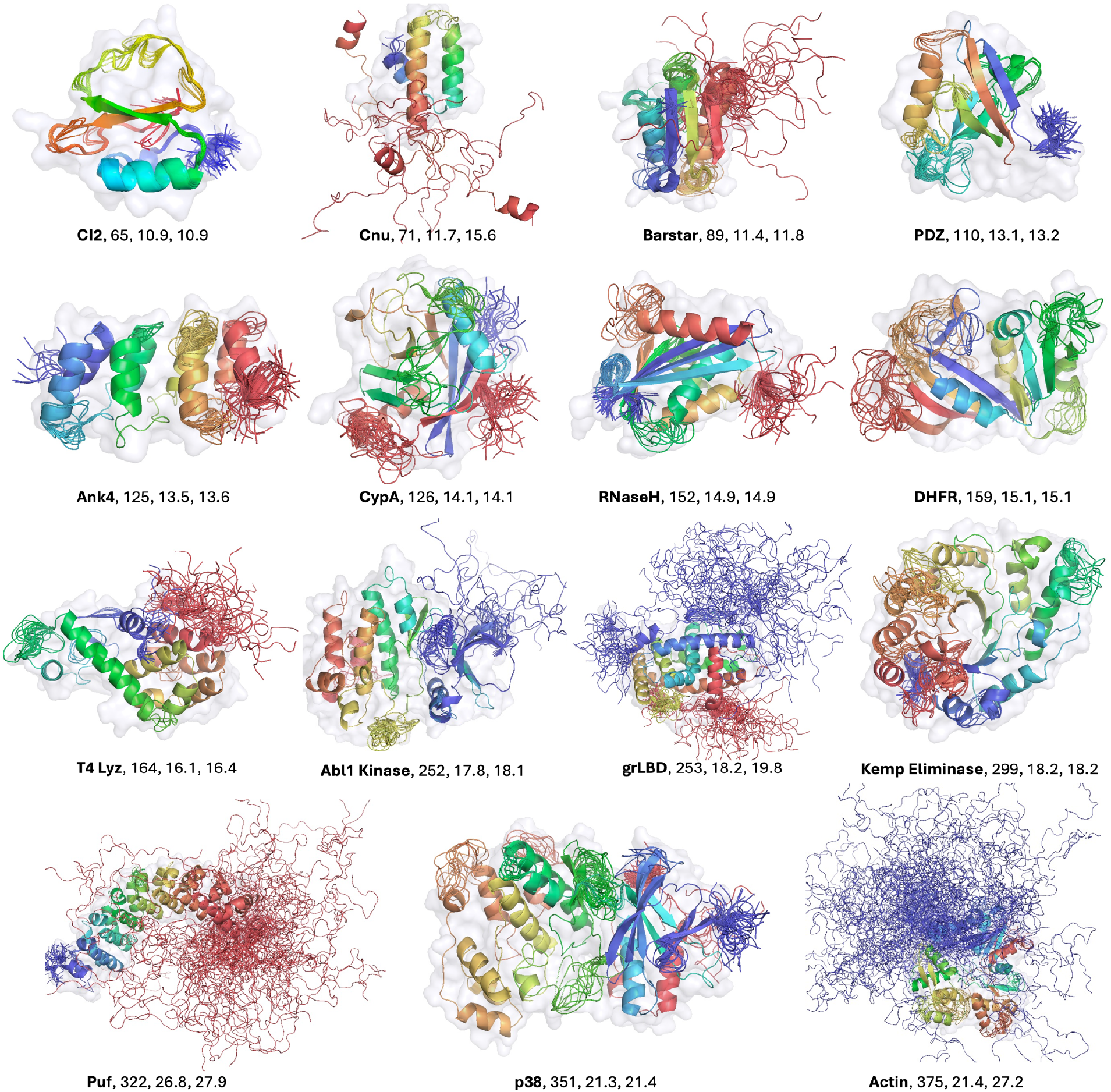
*Hashi*-derived ensembles for a small database of proteins with protein lengths spanning 65-375 residues (Table S1). The ensembles are color coded in the spectral scale with the N- and C-termini in blue and red, respectively. The background surface representation corresponds to the PDB structure. The text below each cartoon represents the following: the protein name, number of residues, radius of gyration of the PDB structure, mean radius of gyration of the ensemble (in Å units).

### Structural excursions driving antibiotic sequestration

Beyond generating native ensembles, this approach can offer mechanistic insight, as illustrated below for the case of an antibiotic sequestration in the protein AlbAS. The antibiotic albicidin sits right into the hydrophobic core of AlbAS spanning both the N-terminal and C-terminal subdomains (NTSD and CTSD, respectively; Figure 4A).^59,61^ We had recently employed the bWSME model to simultaneously reproduce both the global unfolding features (from scanning calorimetry experiments) and local stability profiles (from HDX-MS) to arrive at an experimentally constrained folding conformational landscape of AlbAS (Figure 4B).^59^ Our predictions point to specific melting and structural opening events in the NTSD that could potentially enable access to the buried binding site.

**Figure 4.**
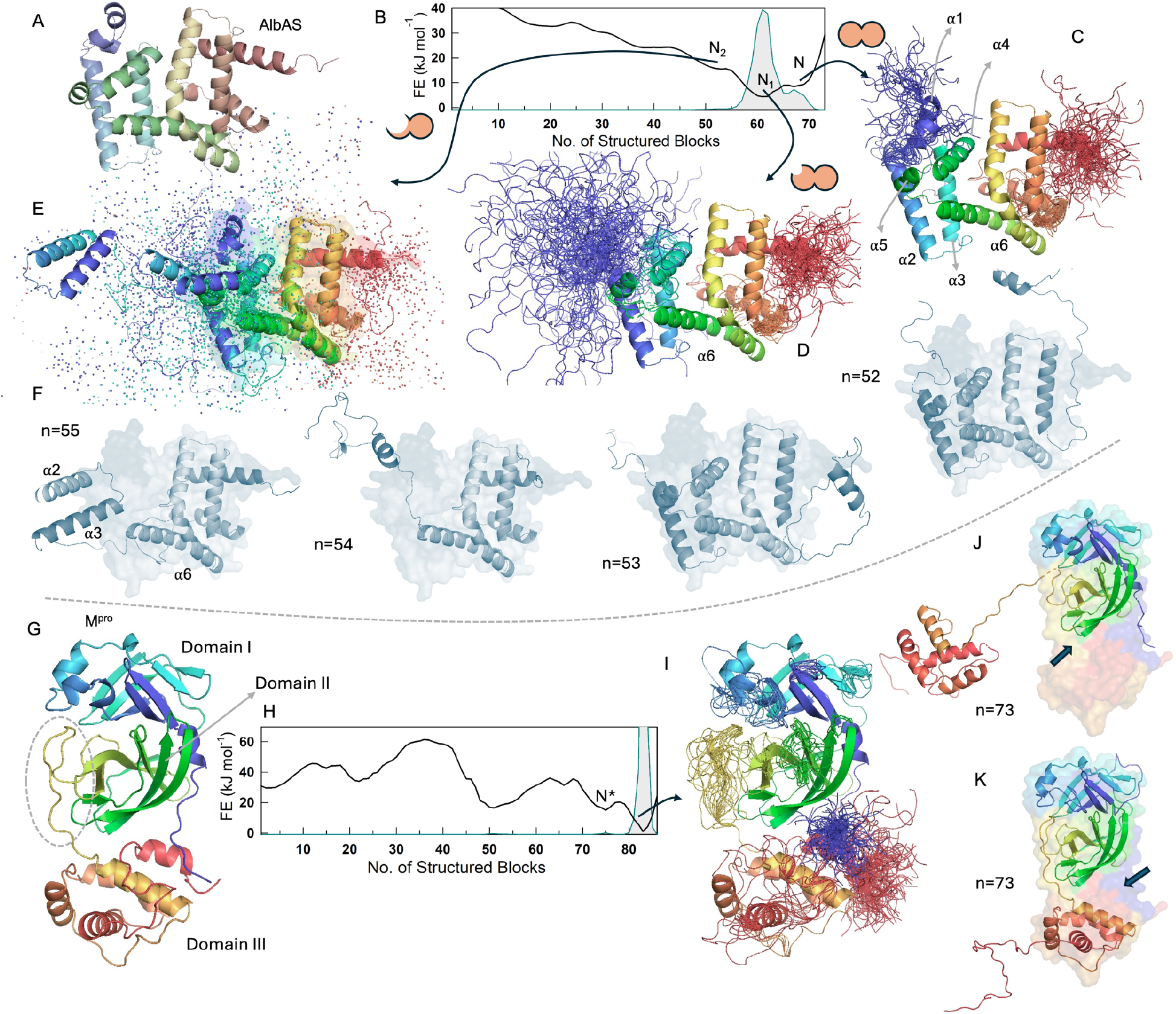
Conformational ensembles of AlbAS and M^pro^. (A) Structure of AlbAS (PDB id: 6ET8). (B) One-dimensional free energy profile of AlbAS and the three macrostates N, N1 and N2. (C, D, E, F) Structural ensembles of the three macrostates colored in the spectral scale from N- to C-terminus (blue to red). The two-circle cartoon depicts the NTSD and CTSD, respectively. Unfolding of the NTSD is represented as a loss of color in the NTSD. Since N2 is a broad ensemble, deviations from the folded structure are represented as dots. Panel F plots representative and most probable structural realizations at the indicated RC values (*n*) as cartoons, while a surface representation of the fully folded state is shown in the background. The parameters are: *macrostate_pools = [65 71; 58 64; 48 55], no_of_microstates = 20, no_of_conformations = 5*. (G, H) Structure of M^pro^ (PDB id: 6M0J) with the long-disordered loop circled (panel G), and the associated free energy profile (panel H). (I, J, K) The native ensemble of the protease (panel I) with specific structural realizations at a RC value of 73 that corresponds to N* (panels J and K). The arrows in panels J and K represent the protein regions exposed during structural excursions. The folded structure is shown in the background. The parameters are: *macrostate_pools = [80 85; 72 77], no_of_microstates = 30, no_of_conformations = 5*.

The *pepval* matrix carrying the AlbAS microstate information was passed through the *Hashi* workflow to generate ensembles corresponding to the macrostate *N*, *N1* and *N2* previously identified. The folded ensemble *N* appears as an excited state along the folding reaction coordinate and is characterized by significant fluctuations in the N- and C-terminal regions (Figure 4C). *N1*, the most populated state in the native ensemble, displays greater disorder in the N-terminal helices compared to the C-terminal helices (Figure 4D). *N2*, the partially structured excited state, displays much more conformational variability primarily in the NTSD (Figure 4E). In fact, the core of the CTSD is fully folded including the *a*6 helix that bridges the two subdomains. The NTSD samples multiple conformational substates driven by disorder in the helices *a*4 and *a*5, while *a*2 and *a*3 remain intact (Figure 4F). The most probable conformation along every reaction coordinate value of the macrostate *N2* provides a vivid picture of the potential structural changes that are consistent with published experimental data.^59^

### Identifying cryptic pockets in a protease

The static structure of SARS-CoV-2 protease M^pro^ is characterized by three domains with a long loop connecting the domains II and III (Figure 4G).^62^ The predicted free energy profile displays a broad native ensemble (Figure 4H),^32^ which arises from substantial loop motions across different protein parts as identified by the *Hashi* workflow (Figure 4I). The excited state at *n* = 73 corresponds to an assortment of microstates in which the domain III is fully undocked (Figure 4J) or partially open, with associated melting of the C-terminal helices (Figure 4K). The undocked conformation of domain III relative to domain I and II has been observed in long time-scale MD simulations.^63^ The *Hashi* pipeline can therefore be employed to identify cryptic pockets^64,65^ in large proteins, in addition to characterizing their unfolding mechanisms.

## Conclusions

The success of the (b)WSME model of protein folding can be attributed to a combination of key model features: a fixed yet physically realistic conformational ensemble composed of strings of *zeros* and *ones*, a simple and tunable energy–entropy framework, and the ability to efficiently compute the total partition function. The *Hashi* workflow presented here establishes a general framework for bridging such string-based representations with a three-dimensional view of the folding landscapes, opening avenues for robust explorations of conformational ensembles and by extension the folding mechanism. This is made possible by the already highly optimized protocol of the RANCH module of EOM that can quickly generate ensembles given the boundaries separating ordered and disordered segments.

The current AI/ML-based generative methods^14–18^ require either large training data sets or enhanced sampling to generate ensembles. *Hashi*, on the other hand, exploits the bWSME model’s approximation of the state space into discrete pre-defined microstates, requiring no training data or intensive computational resources. This allows *Hashi* to generate 3D ensembles in seconds to minutes from a single static structure for a range of monomeric protein systems, irrespective of the degree of disorder, on a standard computer. Importantly, the statistical weights of the microstates from the bWSME model can be directly recalibrated against experimental data before ensemble generation, offering an experimentally derived alternative to the data-driven emulators. *Hashi* is therefore well suited for rapid hypothesis generation and providing physically realistic starting ensembles for all-atom MD simulations, rather than as a replacement for methods offering atomic-level accuracy. Taken together with the tunability intrinsic to the WSME model, the *Hashi* workflow not only bridges the 1D strings with 3D ensembles but also complements both generative and conventional molecular dynamics methods.

## Acknowledgements

The authors thank Alessandra del Giudice (Sapienza University of Rome, Italy) for introducing the RANCH module to us. The authors acknowledge the High Performance Computing Environment (HPCE) facility, IIT Madras (Chennai, India), for providing the computational facilities.

## Competing Financial Interests

The authors declare no competing financial interests.

## Data and Software Availability

The bWSME model code, step-by-step procedure for integrating the output with *Hashi* and analysis functions with representative examples can be found at https://github.com/AthiNaganathan/Hashi.

## TOC Graphic

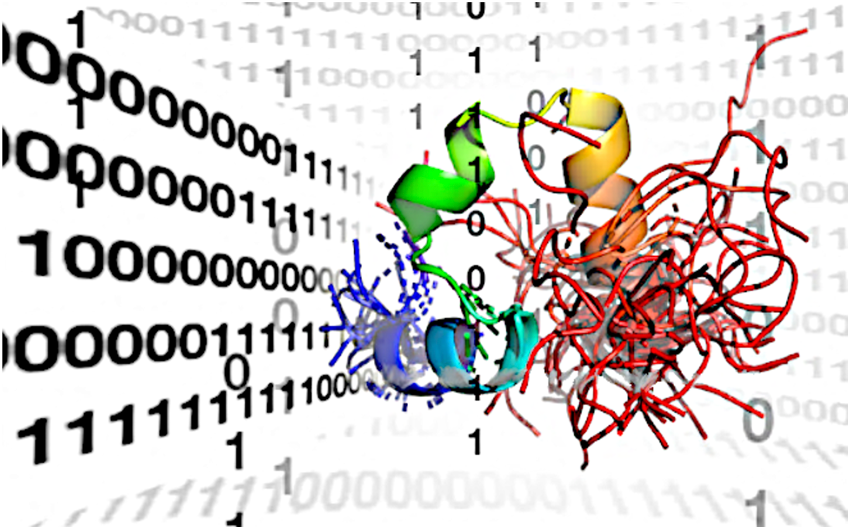

## Notes

https://github.com/AthiNaganathan/Hashi

### Competing Interest Statement

The authors have declared no competing interest.

### Summary of Updates

Following reviewer's comments, an extra figure has been added to the main text including discussions. A supporting file is also included providing additional information.

## REFERENCES

(1) Tokuriki, N.; Tawfik, D. S. Stability Effects of Mutations and Protein Evolvability. Curr. Opin. Struct. Biol. 2009, 19 (5), 596–604. 10.1016/j.sbi.2009.08.003.

(2) Petrovic, D.; Risso, V. A.; Kamerlin, S. C. L.; Sanchez-Ruiz, J. M. Conformational Dynamics and Enzyme Evolution. J. R. Soc. Interface 2018, 15 (144), 20180330. 10.1098/rsif.2018.0330.

(3) Yabukarski, F.; Doukov, T.; Pinney, M. M.; Biel, J. T.; Fraser, J. S.; Herschlag, D. Ensemble-Function Relationships to Dissect Mechanisms of Enzyme Catalysis. Sci. Adv. 2022, 8 (41). 10.1126/sciadv.abn7738.

(4) Henzler-Wildman, K.; Kern, D. Dynamic Personalities of Proteins. Nature 2007, 450 (7172), 964–972. 10.1038/nature06522.

(5) Rueda, M.; Ferrer-Costa, C.; Meyer, T.; Perez, A.; Camps, J.; Hospital, A.; Gelpi, J. L.; Orozco, M. A Consensus View of Protein Dynamics. Proc. Natl. Acad. Sci. U.S.A. 2007, 104 (3), 796–801. 10.1073/pnas.0605534104.

(6) Best, R. B. Atomistic Molecular Simulations of Protein Folding. Curr. Opin. Struct. Biol. 2012, 22, 52–61. 10.1016/j.sbi.2011.12.001.

(7) Piana, S.; Klepeis, J. L.; Shaw, D. E. Assessing the Accuracy of Physical Models Used in Protein-Folding Simulations: Quantitative Evidence from Long Molecular Dynamics Simulations. Curr. Opin. Struct. Biol. 2014, 24, 98–105. 10.1016/j.sbi.2013.12.006.

(8) Chodera, J. D.; Noe, F. Markov State Models of Biomolecular Conformational Dynamics. Curr. Opin. Struct. Biol. 2014, 25, 135–144. 10.1016/j.sbi.2014.04.002.

(9) Levy, Y.; Cho, S. S.; Onuchic, J. N.; Wolynes, P. G. A Survey of Flexible Protein Binding Mechanisms and Their Transition States Using Native Topology Based Energy Landscapes. J. Mol. Biol. 2005, 346, 1121–1145. 10.1016/j.jmb.2004.12.021.

(10) Hyeon, C.; Thirumalai, D. Capturing the Essence of Folding and Functions of Biomolecules Using Coarse-Grained Models. Nat. Commun. 2011, 2, 487. 10.1038/ncomms1481.

(11) Leone, V.; Marinelli, F.; Carloni, P.; Parrinello, M. Targeting Biomolecular Flexibility with Metadynamics. Curr. Opin. Struct. Biol. 2010, 20, 148–154. 10.1016/j.sbi.2010.01.011.

(12) Luitz, M.; Bomblies, R.; Ostermeir, K.; Zacharias, M. Exploring Biomolecular Dynamics and Interactions Using Advanced Sampling Methods. J. Phys. Condens. Matter 2015, 27 (32), 323101. 10.1088/0953-8984/27/32/323101.

(13) Mehdi, S.; Smith, Z.; Herron, L.; Zou, Z.; Tiwary, P. Enhanced Sampling with Machine Learning. Ann. Rev. Phys. Chem. 2024, 75 (1), 347–370. 10.1146/annurev-physchem-083122-125941.

(14) Noé, F.; Olsson, S.; Köhler, J.; Wu, H. Boltzmann Generators: Sampling Equilibrium States of Many-Body Systems with Deep Learning. Science (1979). 2019, 365 (6457). 10.1126/science.aaw1147.

(15) Vani, B. P.; Aranganathan, A.; Wang, D.; Tiwary, P. AlphaFold2-RAVE: From Sequence to Boltzmann Ranking. J. Chem. Theory Comput. 2023, 19 (14), 4351–4354. 10.1021/acs.jctc.3c00290.

(16) Lewis, S.; Hempel, T.; Jiménez-Luna, J.; Gastegger, M.; Xie, Y.; Foong, A. Y. K.; Satorras, V. G.; Abdin, O.; Veeling, B. S.; Zaporozhets, I.; Chen, Y.; Yang, S.; Foster, A. E.; Schneuing, A.; Nigam, J.; Barbero, F.; Stimper, V.; Campbell, A.; Yim, J.; Lienen, M.; Shi, Y.; Zheng, S.; Schulz, H.; Munir, U.; Sordillo, R.; Tomioka, R.; Clementi, C.; Noé, F. Scalable Emulation of Protein Equilibrium Ensembles with Generative Deep Learning. Science (1979). 2025, 389 (6761). 10.1126/science.adv9817.

(17) Charron, N. E.; Bonneau, K.; Pasos-Trejo, A. S.; Guljas, A.; Chen, Y.; Musil, F.; Venturin, J.; Gusew, D.; Zaporozhets, I.; Krämer, A.; Templeton, C.; Kelkar, A.; Durumeric, A. E. P.; Olsson, S.; Pérez, A.; Majewski, M.; Husic, B. E.; Patel, A.; De Fabritiis, G.; Noé, F.; Clementi, C. Navigating Protein Landscapes with a Machine-Learned Transferable Coarse-Grained Model. Nat. Chem. 2025, 17 (8), 1284–1292. 10.1038/s41557-025-01874-0.

18. (18) Olsson, S. Generative Molecular Dynamics. Curr. Opin. Struct. Biol. 2026, 96, 103213. 10.1016/j.sbi.2025.103213.

(19) Taketomi, H.; Ueda, Y.; Go, N. Studies on Protein Folding, Unfolding and Fluctuations by Computer-Simulation .1. Effect of Specific Amino-Acid Sequence Represented by Specific Inter-Unit Interactions. Inter. J. Prot. Pep. Res. 1975, 7 (6), 445–459. 10.1111/j.1399-3011.1975.tb02465.x.

(20) Wako, H.; Saito, N. Statistical Mechanical Theory of Protein Conformation .2. Folding Pathway for Protein. J. Phys. Soc. Japan 1978, 44 (6), 1939–1945. 10.1143/JPSJ.44.1939.

(21) Muñoz, V.; Eaton, W. A. A Simple Model for Calculating the Kinetics of Protein Folding from Three-Dimensional Structures. Proc. Natl. Acad. Sci. U.S.A. 1999, 96 (20), 11311–11316. 10.1073/pnas.96.20.11311.

(22) Gopi, S.; Aranganathan, A.; Naganathan, A. N. Thermodynamics and Folding Landscapes of Large Proteins from a Statistical Mechanical Model. Curr. Res. Struct. Biol. 2019, 1, 6–12. 10.1016/j.crstbi.2019.10.002.

(23) Bruscolini, P.; Naganathan, A. N. Quantitative Prediction of Protein Folding Behaviors from a Simple Statistical Model. J. Am. Chem. Soc. 2011, 133 (14), 5372–5379. 10.1021/ja110884m.

(24) Naganathan, A. N. Predictions from an Ising-like Statistical Mechanical Model on the Dynamic and Thermodynamic Effects of Protein Surface Electrostatics. J. Chem. Theory Comput. 2012, 8 (11), 4646–4656. 10.1021/ct300676w.

(25) Rajasekaran, N.; Gopi, S.; Narayan, A.; Naganathan, A. N. Quantifying Protein Disorder through Measures of Excess Conformational Entropy. J. Phys. Chem. B 2016, 120, 4341– 4350. 10.1021/acs.jpcb.6b00658.

(26) Ooka, K.; Arai, M. Accurate Prediction of Protein Folding Mechanisms by Simple Structure-Based Statistical Mechanical Models. Nat. Commun. 2023, 14 (1), 6338. 10.1038/s41467-023-41664-1.

(27) Itoh, K.; Sasai, M. Flexibly Varying Folding Mechanism of a Nearly Symmetrical Protein: B Domain of Protein A. Proc. Natl. Acad. Sci. U S A 2006, 103 (19), 7298–7303. 10.1073/pnas.0510324103.

(28) Inanami, T.; Terada, T. P.; Sasai, M. Folding Pathway of a Multidomain Protein Depends on Its Topology of Domain Connectivity. Proc. Natl. Acad. Sci. U. S. A. 2014, 111 (45), 15969–15974. 10.1073/pnas.1406244111.

(29) Narayan, A.; Campos, L. A.; Bhatia, S.; Fushman, D.; Naganathan, A. N. Graded Structural Polymorphism in a Bacterial Thermosensor Protein. J. Am. Chem. Soc. 2017, 139, 792–802. 10.1021/jacs.6b10608.

(30) Sivanandan, S.; Naganathan, A. N. A Disorder-Induced Domino-Like Destabilization Mechanism Governs the Folding and Functional Dynamics of the Repeat Protein IκBα. PLOS Comput. Biol. 2013, 9, e1003403. 10.1371/journal.pcbi.1003403.

(31) Ooka, K.; Liu, R.; Arai, M. The Wako-Saitô-Muñoz-Eaton Model for Predicting Protein Folding and Dynamics. Molecules 2022, 27 (14), 4460. 10.3390/molecules27144460.

(32) Naganathan, A. N.; Dani, R.; Gopi, S.; Aranganathan, A.; Narayan, A. Folding Intermediates, Heterogeneous Native Ensembles and Protein Function. J. Mol. Biol. 2021, 433 (24), 167325. 10.1016/j.jmb.2021.167325.

(33) Dani, R.; Kurbah, I.; Chaurasiya, D. K.; Nath, R.; Krishnan, A.; Fushman, D.; Naganathan, A. N. Selective Control of Ligand Binding through a Distal Mutation That Alters the Protein Native Ensemble. Proc. Natl. Acad. Sci. USA 2026, 123 (8). 10.1073/pnas.2519658123.

(34) Naganathan, A. N. A Rapid, Ensemble and Free Energy Based Method for Engineering Protein Stabilities. J. Phys. Chem. B 2013, 117 (17), 4956–4964. 10.1021/jp401588x.

(35) Kannan, A.; Naganathan, A. N. Ensemble Origins and Distance-Dependence of Long-Range Mutational Effects in Proteins. iScience 2022, 25 (10), 105181. 10.1016/j.isci.2022.105181.

(36) Anantakrishnan, S.; Naganathan, A. N. Thermodynamic Architecture and Conformational Plasticity of GPCRs. Nat. Commun. 2023, 14 (1), 128. 10.1038/s41467-023-35790-z.

(37) Naganathan, A. N.; Kannan, A. A Hierarchy of Coupling Free Energies Underlie the Thermodynamic and Functional Architecture of Protein Structures. Curr. Res. Struct. Biol. 2021, 3, 257–267. 10.1016/j.crstbi.2021.09.003.

(38) Chaurasiya, D. K.; Naganathan, A. N. Diverse Conformational Ensembles Define the Shared Folding-Allosteric Landscapes of Protein Kinases. Biophys. J. 2026, 125 (2), 557–571. 10.1016/j.bpj.2025.09.035.

(39) Henry, E. R.; Best, R. B.; Eaton, W. A. Comparing a Simple Theoretical Model for Protein Folding with All-Atom Molecular Dynamics Simulations. Proc. Natl. Acad. Sci. U.S.A. 2013, 110 (44), 17880–17885. 10.1073/pnas.1317105110.

(40) Gopi, S.; Singh, A.; Suresh, S.; Paul, S.; Ranu, S.; Naganathan, A. N. Toward a Quantitative Description of Microscopic Pathway Heterogeneity in Protein Folding. Phys. Chem. Chem. Phys. 2017, 19 (31), 20891–20903. 10.1039/c7cp03011h.

(41) Kannan, A.; Chaurasiya, D. K.; Naganathan, A. N. Conflicting Interfacial Electrostatic Interactions as a Design Principle to Modulate Long-Range Interdomain Communication. ACS Bio. Med. Chem. Au 2024, 4 (1), 53–67. 10.1021/acsbiomedchemau.3c00047.

(42) Bruscolini, P.; Pelizzola, A. Exact Solution of the Muñoz-Eaton Model for Protein Folding. Phys. Rev. Lett. 2002, 88 (25), 258101. 10.1103/physrevlett.88.258101.

(43) Henry, E. R.; Eaton, W. A. Combinatorial Modeling of Protein Folding Kinetics: Free Energy Profiles and Rates. Chem. Phys. 2004, 307, 163–185. 10.1016/j.chemphys.2004.06.064.

(44) Bernado, P.; Mylonas, E.; Petoukhov, M. V; Blackledge, M.; Svergun, D. I. Structural Characterization of Flexible Proteins Using Small-Angle X-Ray Scattering. J. Am. Chem. Soc. 2007, 129 (17), 5656–5664. 10.1021/ja069124n.

(45) Tria, G.; Mertens, H. D. T.; Kachala, M.; Svergun, D. I. Advanced Ensemble Modelling of Flexible Macromolecules Using X-Ray Solution Scattering. IUCrJ 2015, 2 (Pt 2), 207–217. 10.1107/S205225251500202X.

(46) Franke, D.; Gräwert, T.; Svergun, D. I. New Features in ATSAS-4, a Program Suite for Small-Angle Scattering Data Analysis. J. Appl. Cryst. 2025, 58 (3), 1027–1033. 10.1107/S1600576725002481.

(47) Rotkiewicz, P.; Skolnick, J. Fast Procedure for Reconstruction of Full-atom Protein Models from Reduced Representations. J. Comput. Chem. 2008, 29 (9), 1460–1465. 10.1002/jcc.20906.

(48) Wang, Q.; Canutescu, A. A.; Dunbrack, R. L. SCWRL and MolIDE: Computer Programs for Side-Chain Conformation Prediction and Homology Modeling. Nat. Protoc. 2008, 3 (12), 1832–1847. 10.1038/nprot.2008.184.

(49) Kubelka, J.; Henry, E. R.; Cellmer, T.; Hofrichter, J.; Eaton, W. A. Chemical, Physical, and Theoretical Kinetics of an Ultrafast Folding Protein. Proc. Natl. Acad. Sci. USA 2008, 105 (48), 18655–18662. 10.1073/pnas.0808600105.

(50) Cellmer, T.; Henry, E. R.; Kubelka, J.; Hofrichter, J.; Eaton, W. A. Relaxation Rate for an Ultrafast Folding Protein Is Independent of Chemical Denaturant Concentration. J. Am. Chem. Soc. 2007, 129 (47), 14564–14565. 10.1021/ja0761939.

(51) Kubelka, J.; Chiu, T. K.; Davies, D. R.; Eaton, W. A.; Hofrichter, J. Sub-Microsecond Protein Folding. J. Mol. Biol. 2006, 359 (3), 546–553. 10.1016/j.jmb.2006.03.034.

(52) Godoy-Ruiz, R.; Henry, E. R.; Kubelka, J.; Hofrichter, J.; Munoz, V.; Sanchez-Ruiz, J. M.; Eaton, W. A. Estimating Free-Energy Barrier Heights for an Ultrafast Folding Protein from Calorimetric and Kinetic Data. J. Phys. Chem. B 2008, 112 (19), 5938–5949. 10.1021/jp0757715.

(53) Bi, Y.; Cho, J. H.; Kim, E. Y.; Shan, B.; Schindelin, H.; Raleigh, D. P. Rational Design, Structural and Thermodynamic Characterization of a Hyperstable Variant of the Villin Headpiece Helical Subdomain. Biochemistry 2007, 46 (25), 7497–7505. 10.1021/bi6026314.

(54) Reiner, A.; Henklein, P.; Kiefhaber, T. An Unlocking/Relocking Barrier in Conformational Fluctuations of Villin Headpiece Subdomain. Proc. Natl. Acad. Sci. U.S.A. 2010, 107 (11), 4955–4960. 10.1073/pnas.0910001107.

(55) Luan, Q.; Clark, P. L. Identification of an On-Pathway Intermediate Illuminates the Kinetic Competition between Protein Folding and Misfolding. Proc. Natl. Acad. Sci. USA 2025, 122 (31). 10.1073/pnas.2425999122.

(56) Dani, R.; Pawloski, W.; Chaurasiya, D. K.; Srilatha, N. S.; Agarwal, S.; Fushman, D.; Naganathan, A. N. Conformational Tuning Shapes the Balance between Functional Promiscuity and Specialization in Paralogous Plasmodium Acyl-CoA Binding Proteins. Biochemistry 2023, 62 (20), 2982–2996. 10.1021/acs.biochem.3c00449.

(57) Natarajan, L.; De Sciscio, M. L.; Nardi, A. N.; Sekhar, A.; Del Giudice, A.; D’Abramo, M.; Naganathan, A. N. A Finely Balanced Order-Disorder Equilibrium Sculpts the Folding-Binding Landscape of an Antibiotic Sequestering Protein. Proc. Nat. Acad. Sci. U.S.A 2024, 121 (20), e2318855121. 10.1073/pnas.2318855121.

(58) Truhlar, S. M. E.; Torpey, J. W.; Komives, E. A. Regions of I Kappa B Alpha That Are Critical for Its Inhibition of NF-Kappa B - DNA Interaction Fold upon Binding to NF-Kappa B. Proc. Natl. Acad. Sci. U.S.A. 2006, 103 (50), 18951–18956. 10.1073/pnas.0605794103.

(59) Natarajan, L.; Loginov, D.; Kadek, A.; Man, P.; Naganathan, A. N. Local and Global Breathing Motions Prime the Access to Buried Binding Site in an Antibiotic-Sequestering Protein. ACS Bio & Med Chem Au 2025, 5 (5), 840–851. 10.1021/acsbiomedchemau.5c00081.

(60) Golla, H.; Kannan, A.; Gopi, S.; Murugan, S.; Perumalsamy, L. R.; Naganathan, A. N. Structural-Energetic Basis for Coupling between Equilibrium Fluctuations and Phosphorylation in a Protein Native Ensemble. ACS Cent. Sci. 2022, 8 (2), 282–293. 10.1021/acscentsci.1c01548.

(61) Rostock, L.; Driller, R.; Grätz, S.; Kerwat, D.; von Eckardstein, L.; Petras, D.; Kunert, M.; Alings, C.; Schmitt, F.-J.; Friedrich, T.; Wahl, M. C.; Loll, B.; Mainz, A.; Süssmuth, R. D. Molecular Insights into Antibiotic Resistance - How a Binding Protein Traps Albicidin. Nat. Commun. 2018, 9 (1), 3095. 10.1038/s41467-018-05551-4.

(62) Jin, Z.; Du, X.; Xu, Y.; Deng, Y.; Liu, M.; Zhao, Y.; Zhang, B.; Li, X.; Zhang, L.; Peng, C.; Duan, Y.; Yu, J.; Wang, L.; Yang, K.; Liu, F.; Jiang, R.; Yang, X.; You, T.; Liu, X.; Yang, X.; Bai, F.; Liu, H.; Liu, X.; Guddat, L. W.; Xu, W.; Xiao, G.; Qin, C.; Shi, Z.; Jiang, H.; Rao, Z.; Yang, H. Structure of M(pro) from SARS-CoV-2 and Discovery of Its Inhibitors. Nature 2020, 582 (7811), 289–293. 10.1038/s41586-020-2223-y.

(63) Zimmerman, M. I.; Porter, J. R.; Ward, M. D.; Singh, S.; Vithani, N.; Meller, A.; Mallimadugula, U. L.; Kuhn, C. E.; Borowsky, J. H.; Wiewiora, R. P.; Hurley, M. F. D.; Harbison, A. M.; Fogarty, C. A.; Coffland, J. E.; Fadda, E.; Voelz, V. A.; Chodera, J. D.; Bowman, G. R. SARS-CoV-2 Simulations Go Exascale to Predict Dramatic Spike Opening and Cryptic Pockets across the Proteome. Nat. Chem. 2021, DOI: 10.1038/s41557-021-00707-0. 10.1038/s41557-021-00707-0.

(64) Bowman, G. R.; Geissler, P. L. Equilibrium Fluctuations of a Single Folded Protein Reveal a Multitude of Potential Cryptic Allosteric Sites. Proc. Natl. Acad. Sci. U. S. A. 2012, 109 (29), 11681–11686. 10.1073/pnas.1209309109.

(65) Zhang, S.; Bowman, G. R. Decrypting Cryptic Pockets with Physics-Based Simulations and Artificial Intelligence. Curr. Opin. Struct. Biol. 2026, 96, 103215. 10.1016/j.sbi.2025.103215.

