## Supporting Information for "*Hashi*: Bridging Statistical Model Derived 1D Microstate Encodings and Protein 3D Structural Ensembles"

AUTHOR INFORMATION

**Corresponding Author**


**Supporting Text**

***Tuning bWSME model parameters to generate free energy profiles*** The code main_WSME.m has options for modifying the input parameters. We describe here the procedure for generating free energy profile for Villin at 310 K. The line numbers 13-23 take in the protein name ('Villin'), block length (a value of 1 is included here, a value of till 6 has been tested), and whether given protein is considered "large" (free energy surfaces for large proteins, >200 residues, are calculated using FesCalc_Block_gen.m instead of FesCalc_Block.m, as detailed below). The pH and distance cutoff to identify vdW interactions (*srcutoff* in Å) are defined in lines 17-18. In addition, the input protein structure (in .pdb and .cif formats), and a text file ('struct.txt', line 16) that carries the output from STRIDE secondary structure assignment tool should be present. The user should ensure that the input structure files contain no missing atoms or residues, and should remove cofactors/water molecules, i.e. no HETATM field should be present. Running this code will generate five files that will be saved in the local folder. Following this, the code 'FesCalc_Block.m' will take in the files generated from the previous code with the line numbers 15 - 19 taking user defined parameters for the entropic penalty for fixing a residue in the native conformation ($\Delta S_{conf}$, included as *DS* in the code), vdW interaction energy per native contact (*ene*), heat capacity change per native contact ($\Delta C_p^{cont}$, *DCp*), temperature ($T$ in K) and ionic strength (*IS* in Molar units). Residue positions that should be assigned larger entropic penalty (coil or glycine residues, *disr*) or lower entropic penalty (proline residues, *ppos*), need not be explicitly defined, as this is automatically generated by the code from the STRIDE output.

A similar protocol is employed for Pertactin (a large 539-residue repeat protein) but with one modification. Given the large number of residues and energy terms, the individual terms in the statistical weights must be calculated in separate windows to avoid numerical blow-up and rounding-off errors; these window-wise terms are then multiplied together. This is implemented in FesCalc_Block_gen.m instead of FesCalc_Block.m. In principle, the former can be used for proteins of any length. Example input parameters for Pertactin are commented out in lines 25-36. The *pepval* matrix and the free energy profile will be generated in *~15 minutes* for a block length of 5 residues.

***Hashi's run time*** *Hashi*'s performance in terms of running time was evaluated for 4 proteins (TipAS, Kemp Eliminase, p38, and Pertactin). To do this, the one-dimensional free energy profile was split into pools consisting of 5 macrostates each. In each pool, the top 10 microstates were chosen, and 10 conformations were generated for each (100 conformations overall per pool of macrostates). Additionally, the generation of conformations per pool was repeated 10 times to account for variations arising from the random-seed initialization of the RANCH module, as well as any background processes in the computer that might have slowed the run time.

In general, it can be seen that conformations can be generated under 20 seconds for most pools irrespective of the degree of order or disorder (Figure S3). Two aberrancies are worth mentioning:

1. Longer run time for Pertactin overall, with few microstates timing out.

   RANCH runs were capped after 2 minutes per microstate. Out of all the proteins tested, only Pertactin showed instances of this time limit actually being encountered. In fact, for the runs carried out, 19 out of the ~140 microstates timed out. Re-running RANCH on the microstates that timed out showed that even with longer time budgets of ~30 min per microstate, conformations for many of these microstates were still not generated.

Analysis of the 19 microstates that timed out revealed that all of them were DSAw/L approximations, and had short, disordered linked regions between the two structured blocks in one of two places (Figure S4A):

a. Residues 231-235 (AGGAV)
b. Residues 525-528 (NGNGQ)

Since DSAw/L constrains the structured blocks to their native positions, it is possible that the short linker length combined with the large conformational flexibility of glycine residues made it difficult to generate proper linker orientations. The unusually longer runs for Pertactin could therefore be a consequence of its repeat nature with multiple short beta-strands followed by short connecting loops.

2. The jump in runtime for p38 for the pool with 56-60 structured blocks (normalized reaction coordinate value of 0.75; ~200 seconds compared to ~5-15 seconds for the rest of the pools)

   As before, it was identified that the jump in overall count was due to 3 microstates that were also in the DSAw/L approximation. Here, the linkers were centred around residues 156-181 (LAVNEDCELKILDFGLARHTDDEMTG; Figure S4B). Note that the conformations for these microstates were generated, and RANCH did not fail or timeout. Taken together with observations in Pertactin, it is likely that RANCH has difficulty efficiently sampling conformations for certain stretches when residues flanking them have restricted conformational mobility.

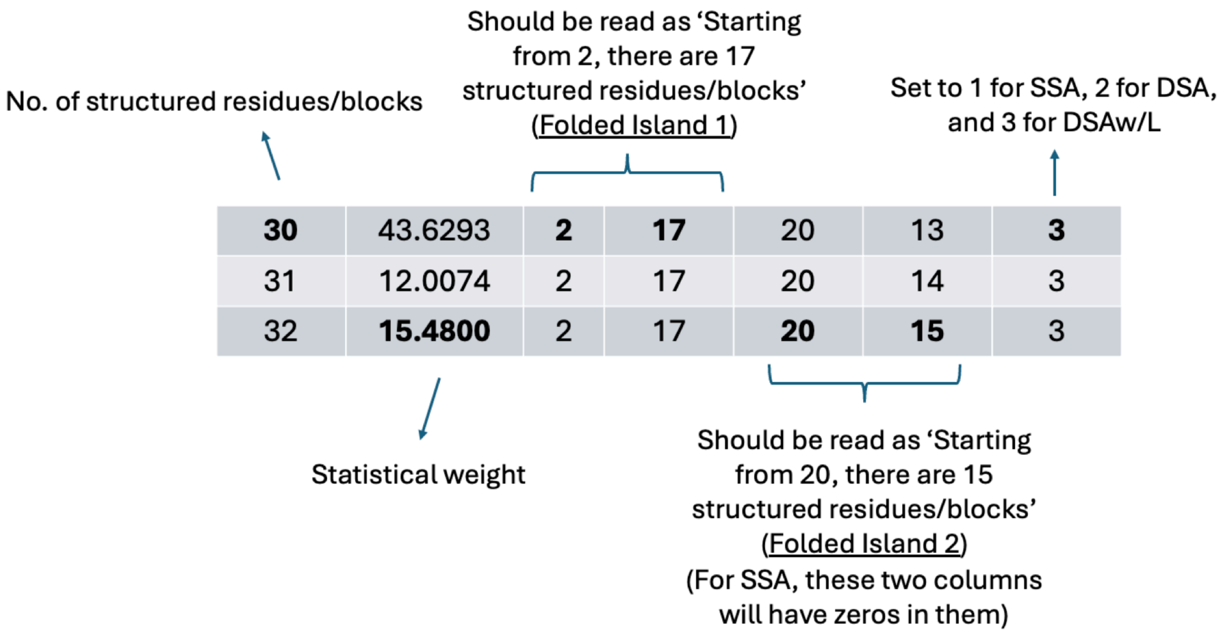


**Figure S1** The contents of the *pepval* matrix – an illustrative example. Every row corresponds to a microstate, while the columns provide information on the total number of residues/blocks that are folded (column 1), the microstate's statistical weight (column 2), structural information (i.e. the regions of the protein that are structured or folded; columns 3-6), and the approximation used (column 7). SSA, DSA and DSAw/L stand for single sequence approximation, double sequence approximation and double sequence approximation with loop, respectively. SSA has a single folded island (a single stretch of *1*s) while DSA and DSAw/L have two folded islands (two stretches of *1*s separated by at least a single unfolded residue/block).

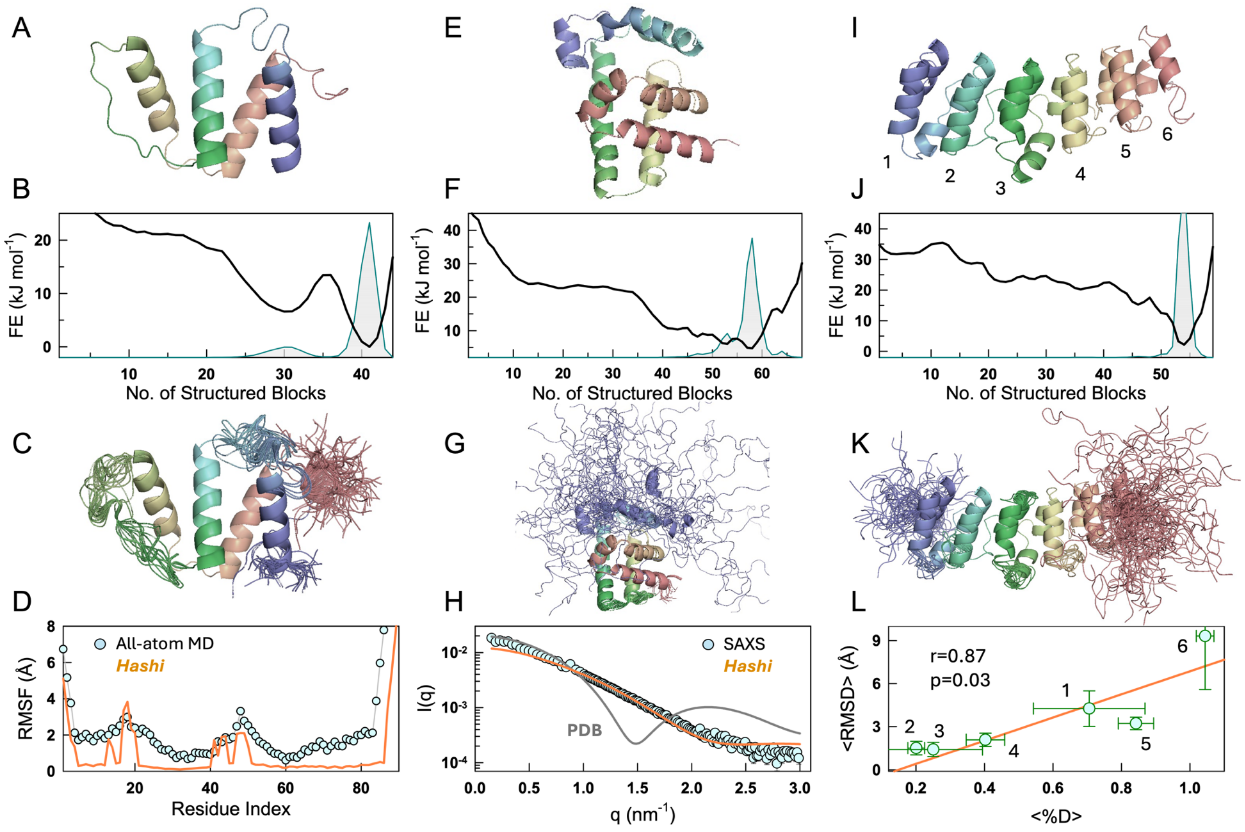


**Figure S2** The first row displays the PDB structures, the second row the bWSME model predicted free energy profiles (black) and the associated probability densities (filled area), the third row the *Hashi*-derived ensembles, and the fourth row the comparisons with experimental data for ACBP (left column; panels A-D), TipAS (middle column; panels E-G) and IκBα (right column; panels I-L). The PDB structures and ensembles are colored in the spectral scale from N- (blue) to C-terminus (red). See Table S1 for PDB files and parameters. (D) All-atom MD derived $C_\alpha$ root-mean-squared fluctuations (circles) versus the same from the *Hashi* ensemble (orange lines) for ACBP. The regions of high flexibility match between the two ensembles. The free-energy profile was predicted after calibrating the model parameters with the experimental heat capacity profile.[1] (H) Experimental SAXS-profile of TipAS (circles) compared to the predicted SAXS profile from the *Hashi* ensemble (orange curve). Note that the free-energy profile was calibrated not with the SAXS data, but with the heat capacity curve.[2] The predicted SAXS profile from the PDB structure 2MBZ is show in dark gray for comparison. (L) The predicted mean $C_\alpha$-RMSD within individual repeats as a function of the percentage deuteration as reported in the experimental study (circles).[3] The orange line is the best fit line with a correlation coefficient of 0.87. Note that the bWSME model was not calibrated with any experimental data for IκBα.

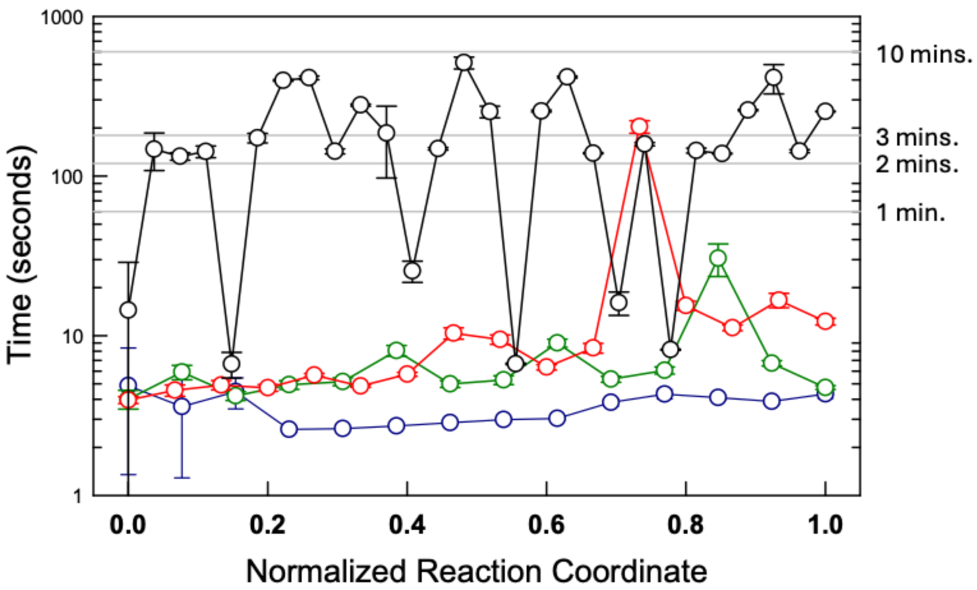


**Figure S3** Mean run times (circles) for generating 10 conformations for the top-ranked 10 microstates in every macrostate pool along the reaction coordinate which is the fraction of structured blocks (the normalized variable is used as the proteins are of different lengths). The error-bars represent the standard deviation in run times from 10 independent runs. The proteins studied are TipAS (143 residues; dark blue), Kemp Eliminase (299 residues; dark green), p38 (351 residues; red) and Pertactin (539 residues; black).

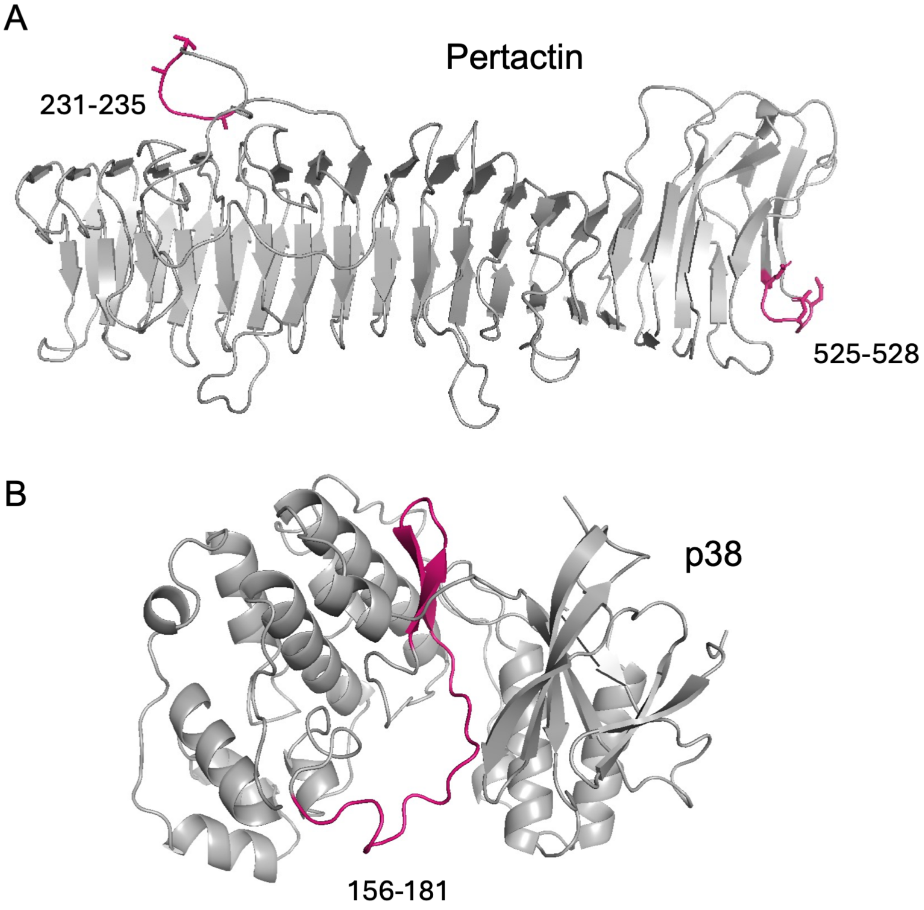


**Figure S4** Structures of Pertactin (panel A) and p38 (panel B). In Pertactin, RANCH failed to generate certain microstates in which the colored stretches were unfolded (i.e., 0s in the bWSME model description). In p38, generating conformations for the stretch containing the long disordered loop (colored) took longer than average. In both cases, the microstates in question belonged to the DSAw/L approximation, in which two folded stretches interact with each other while connected via an unfolded region.

**Table S1** Database of proteins studied.*

| S. No. | Protein | PDB id | Number of Residues (N) | Block Length | Number of Blocks ($n_b$) | Number of Microstates | vdW Interaction Energy (in J mol$^{-1}$) |
|---|---|---|---|---|---|---|---|
| 1 | **Villin** | 1VII | **35** | 1 | 35 | **118,440** | -93.0 |
| 2 | **Pertactin** | 1DAB | **539** | 5 | 137 | **28,935,633** | -79.0 |
| 3 | **pfACBP[1]**** | 1HBK | **90** | 2 | 44 | **289,122** | -67.5 |
| 4 | **TipAS[2]**** | 2MBZ | **143** | 2 | 68 | **1,702,915** | -77.3 |
| 5 | **IκBα** | 1NFI | **213** | 3 | 59 | **936,695** | -80.0 |
| 6 | **AlbAS[4]**** | 6ET8 | **221** | 3 | 73 | **2,258,647** | -37.0 |
| 7 | **CI2** | 2CI2 | **65** | 1 | 65 | **1,372,804** | -88.0 |
| 8 | **Cnu[5]***** | - | **71** | 2 | 33 | **89,687** | -78.5 |
| 9 | **Barstar** | 1BTA | **89** | 2 | 41 | **220,554** | -78.0 |
| 10 | **PDZ** | 1BE9 | **110** | 2 | 50 | **485,266** | -88.0 |
| 11 | **Ank4******** | 1OT8 | **125** | 2 | 60 | **1,007,723** | -91.0 |
| 12 | **CypA** | 1M9F | **165** | 3 | 55 | **721,951** | -79.5 |
| 13 | **RNase H** | 1F21 | **152** | 3 | 50 | **490,450** | -78.0 |
| 14 | **DHFR** | 7DFR | **159** | 3 | 53 | **614,138** | -81.0 |
| 15 | **T4 Lysozyme** | 2LCB | **164** | 3 | 57 | **830,997** | -76.0 |
| 16 | **Abl1 Kinase** | 6XR6 | **252** | 3 | 85 | **4,190,330** | -75.4 |
| 17 | **grLBD** | 1M2Z | **253** | 4 | 65 | **1,420,827** | -71.0 |
| 18 | **HG3.17 Kemp Eliminase** | 4BS0 | **299** | 5 | 71 | **2,002,326** | -73.0 |
| 19 | **Puf** | 1IB2 | **322** | 5 | 86 | **4,384,627** | -67.0 |
| 20 | **p38** | 1W82 | **351** | 5 | 79 | **3,132,726** | -74.5 |
| 21 | **Actin** | 6C1D | **375** | 3 | 127 | **20,977,574** | -77.0 |
| 22 | **SARS-CoV-2 $M^{Pro}$** | 6M0J | **194** | 3 | 64 | **1,310,910** | -81.5 |

* For proteins 7-22 the entire conformational ensemble (i.e. all microstates within the *pepval* matrix) was rank ordered based on their statistical weights. Following this, the top 20 microstates were considered and 5 conformations per microstate was generated using the *Hashi* workflow.

** The references provide additional detail on bWSME model calibration.

*** Full-length structure modeled using AlphaFold3.[6]

**** Only residues 51-175 that constitute the first four Ankyrin repeats were considered.

### Supporting References


(1) Dani, R.; Pawloski, W.; Chaurasiya, D. K.; Srilatha, N. S.; Agarwal, S.; Fushman, D.; Naganathan, A. N. Conformational Tuning Shapes the Balance between Functional Promiscuity and Specialization in Paralogous Plasmodium Acyl-CoA Binding Proteins. *Biochemistry* **2023**, *62* (20), 2982–2996. https://doi.org/10.1021/acs.biochem.3c00449.

(2) Natarajan, L.; De Sciscio, M. L.; Nardi, A. N.; Sekhar, A.; Del Giudice, A.; D'Abramo, M.; Naganathan, A. N. A Finely Balanced Order-Disorder Equilibrium Sculpts the Folding-Binding Landscape of an Antibiotic Sequestering Protein. *Proc. Nat. Acad. Sci. U.S.A* **2024**, *121* (20), e2318855121. https://doi.org/10.1073/pnas.2318855121.

(3) Truhlar, S. M. E.; Torpey, J. W.; Komives, E. A. Regions of I Kappa B Alpha That Are Critical for Its Inhibition of NF-Kappa B - DNA Interaction Fold upon Binding to NF-Kappa B. *Proc. Natl. Acad. Sci. U.S.A.* **2006**, *103* (50), 18951–18956. https://doi.org/10.1073/pnas.0605794103.

(4) Natarajan, L.; Loginov, D.; Kadek, A.; Man, P.; Naganathan, A. N. Local and Global Breathing Motions Prime the Access to Buried Binding Site in an Antibiotic-Sequestering Protein. *ACS Bio & Med Chem Au* **2025**, *5* (5), 840–851. https://doi.org/10.1021/acsbiomedchemau.5c00081.

(5) Narayan, A.; Campos, L. A.; Bhatia, S.; Fushman, D.; Naganathan, A. N. Graded Structural Polymorphism in a Bacterial Thermosensor Protein. *J. Am. Chem. Soc.* **2017**, *139*, 792–802. https://doi.org/10.1021/jacs.6b10608.

(6) Abramson, J.; Adler, J.; Dunger, J.; Evans, R.; Green, T.; Pritzel, A.; Ronneberger, O.; Willmore, L.; Ballard, A. J.; Bambrick, J.; Bodenstein, S. W.; Evans, D. A.; Hung, C.-C.; O'Neill, M.; Reiman, D.; Tunyasuvunakool, K.; Wu, Z.; Žemgulytė, A.; Arvaniti, E.; Beattie, C.; Bertolli, O.; Bridgland, A.; Cherepanov, A.; Congreve, M.; Cowen-Rivers, A. I.; Cowie, A.; Figurnov, M.; Fuchs, F. B.; Gladman, H.; Jain, R.; Khan, Y. A.; Low, C. M. R.; Perlin, K.; Potapenko, A.; Savy, P.; Singh, S.; Stecula, A.; Thillaisundaram, A.; Tong, C.; Yakneen, S.; Zhong, E. D.; Zielinski, M.; Žídek, A.; Bapst, V.; Kohli, P.; Jaderberg, M.; Hassabis, D.; Jumper, J. M. Accurate Structure Prediction of Biomolecular Interactions with AlphaFold 3. *Nature* **2024**, *630* (8016), 493–500. https://doi.org/10.1038/s41586-024-07487-w.
